# CD8+ lymphocytes modulate Zika virus dynamics and tissue dissemination and orchestrate antiviral immunity

**DOI:** 10.1101/475418

**Authors:** Blake Schouest, Marissa Fahlberg, Elizabeth A. Scheef, Matthew J. Ward, Kyra Headrick, Dawn M. Szeltner, Robert V. Blair, Margaret H. Gilbert, Lara A. Doyle-Meyers, Victoria W. Danner, Myrna C. Bonaldo, Dawn M. Wesson, Antonito T. Panganiban, Nicholas J. Maness

**Author notes:** Corresponding author, Nicholas J Maness.

## Abstract

CD8+ lymphocytes are critically important in the control of viral infections, but their roles in acute Zika virus (ZIKV) infection remain incompletely explored in a model sufficiently similar to humans immunologically. Here, we use CD8+ lymphocyte depletion to dissect acute immune responses in adult male rhesus and cynomolgus macaques infected with ZIKV. CD8 depletion delayed serum viremia and dysregulated patterns of innate immune cell homing and monocyte-driven transcriptional responses in the blood. CD8-depleted macaques also showed evidence of compensatory adaptive immune responses, with elevated Th1 activity and persistence of neutralizing antibodies beyond the clearance of serum viremia. The absence of CD8+ lymphocytes increased viral burdens in lymphatic tissues, semen, and cerebrospinal fluid, and neural lesions were also evident in both CD8-depleted rhesus macaques. Together, these data support a role for CD8+ lymphocytes in the control of ZIKV dissemination and in maintaining immune regulation during acute infection of nonhuman primates.

## Introduction

ZIKV has been a known pathogen for over half a century (Dick et al., 1952), but severe manifestations of the disease were not directly associated with the virus for most of its history. Although recent outbreaks of ZIKV in the Western hemisphere are notorious for neurological complications including congenital Zika syndrome (CZS) and Guillain-Barré syndrome (GBS), most cases remain asymptomatic, and when symptoms arise, they are usually mild and self-limiting (Plourde and Bloch, 2016). Differential immune responses to ZIKV infection may dictate the severity of accompanying diseases and underlie clinical outcomes.

As the immunological correlates of protection against ZIKV are beginning to be explored, the CD8 T cell response is emerging as an important mediator of viral control, as is true with other flaviviruses (Slon Campos et al., 2018). Studies in mice have identified CD8 T cell responses to ZIKV infection, but the induction of ZIKV-associated pathology in these models requires deficiency in type-I interferon (IFN) signaling (Elong Ngono et al., 2017; Huang et al., 2017), which is not representative of natural ZIKV infection in humans. This is perhaps an unavoidable caveat, as ZIKV is incapable of antagonizing type-I IFN signaling in mice as it does in humans due to a lack of recognition of murine STAT2 by ZIKV NS5 (Grant et al., 2016). Disrupting IFN-I signaling, either genetically or through antibody blockade is, therefore, necessary to recapitulate ZIKV neurotropism in mouse models. Importantly, these studies have described dual protective and deleterious effects of CD8 T cell responses in ZIKV infected mice. While CD8+ lymphocyte infiltration appears to reduce viral burdens in the brain, spinal cord and lymphatic tissue (Elong Ngono et al., 2017; Huang et al., 2017), under certain circumstances, CD8+ influx may also promote immunopathology, evidenced by neural damage and paralysis (Jurado et al., 2018). However, these findings have yet to be replicated in a model sufficiently similar to humans genetically and immunologically. Given the recent development of rhesus (Coffey et al., 2017; Dudley et al., 2016; Dudley et al., 2018; Magnani et al., 2018a; Osuna et al., 2016a) and cynomolgus (Koide et al., 2016; Osuna et al., 2016a) macaque models of ZIKV infection, we sought to explore the role of CD8+ lymphocytes in acute ZIKV infection by way of CD8+ lymphocyte depletion. CD8+ depletion is a well-established immune manipulation in nonhuman primates (Schmitz et al., 1999) and is thus a plausible approach to gauge how the absence of CD8+ cells impacts acute viremia and potentially modulates adaptive responses.

In the present study, we infected four adult male rhesus macaques and five adult male cynomolgus macaques with a minimally passaged Brazilian ZIKV strain. Prior to infection, two animals of each species were depleted of CD8+ lymphocytes, including CD8 T cells and natural killer (NK) cells. The absence of CD8+ lymphocytes resulted in striking virus-response patterns that were not evident in nondepleted macaques, including delayed serum viremia, enhanced viral dissemination to peripheral tissues, and global repression of antiviral gene transcription. CD8-depleted rhesus macaques also manifested brainstem lesions that were characterized by increased inflammation. Finally, the absence of CD8+ lymphocytes appeared to alter patterns of monocyte expansion and activation and induce compensatory adaptive responses, characterized by enhanced Th1 phenotypes and prolonged neutralizing antibody production.

## Methods

### Animal experiments

The four adult male Indian origin rhesus macaques (*Macaca mulatta*) and five adult male cynomolgus macaques (*Macaca fascicularis*) utilized in this study were housed at the Tulane National Primate Research Center (TNPRC). The TNPRC is fully accredited by AAALAC International (Association for the Assessment and Accreditation of Laboratory Animal Care), Animal Welfare Assurance No. A3180-01. Animals were cared for in accordance with the NRC Guide for the Care and Use of Laboratory Animals and the Animal Welfare Act Animal experiments were approved by the Institutional Animal Care and Use Committee of Tulane University (protocol P0367).

Two rhesus macaques (R25671 and R64357) and two cynomolgus macaques (C78777 and C18942) were depleted of CD8+ lymphocytes by administration of the anti-CD8α antibody MT807R1 (NHP Reagent Resource; https://www.nhpreagents.org) (Schmitz et al., 1999). The initial subcutaneous administration of 10 mg/kg at 14 days pre-infection was followed by three intravenous administrations of 5 mg/kg at 11, 7, and 5 days pre-infection, as per the distributor’s protocol. C84545 was treated with the irrelevant control antibody anti-desmipramine (NHP Reagent Resource; https://www.nhpreagents.org) at the same dosages and time intervals pre-infection. All animals were subcutaneously infected with 10^4^ PFU of a Brazilian ZIKV isolate (Bonaldo et al., 2016) at 0 days post-infection (dpi) (Fig. 1a). As part of a previous study, C46456 (nondepleted cynomolgus macaque) was splenectomized 9 months and 19 days prior to inoculation with ZIKV. For data comparison, we included viral loads and complete blood count (CBC) data from a previous cohort of 4 non-pregnant female rhesus macaques (R32835, R24547, R25508, R22624) that were similarly infected with the same dose of the same Brazilian ZIKV isolate that was used in this study (Supplementary Figs. 2a-e).

**Figure 1.**
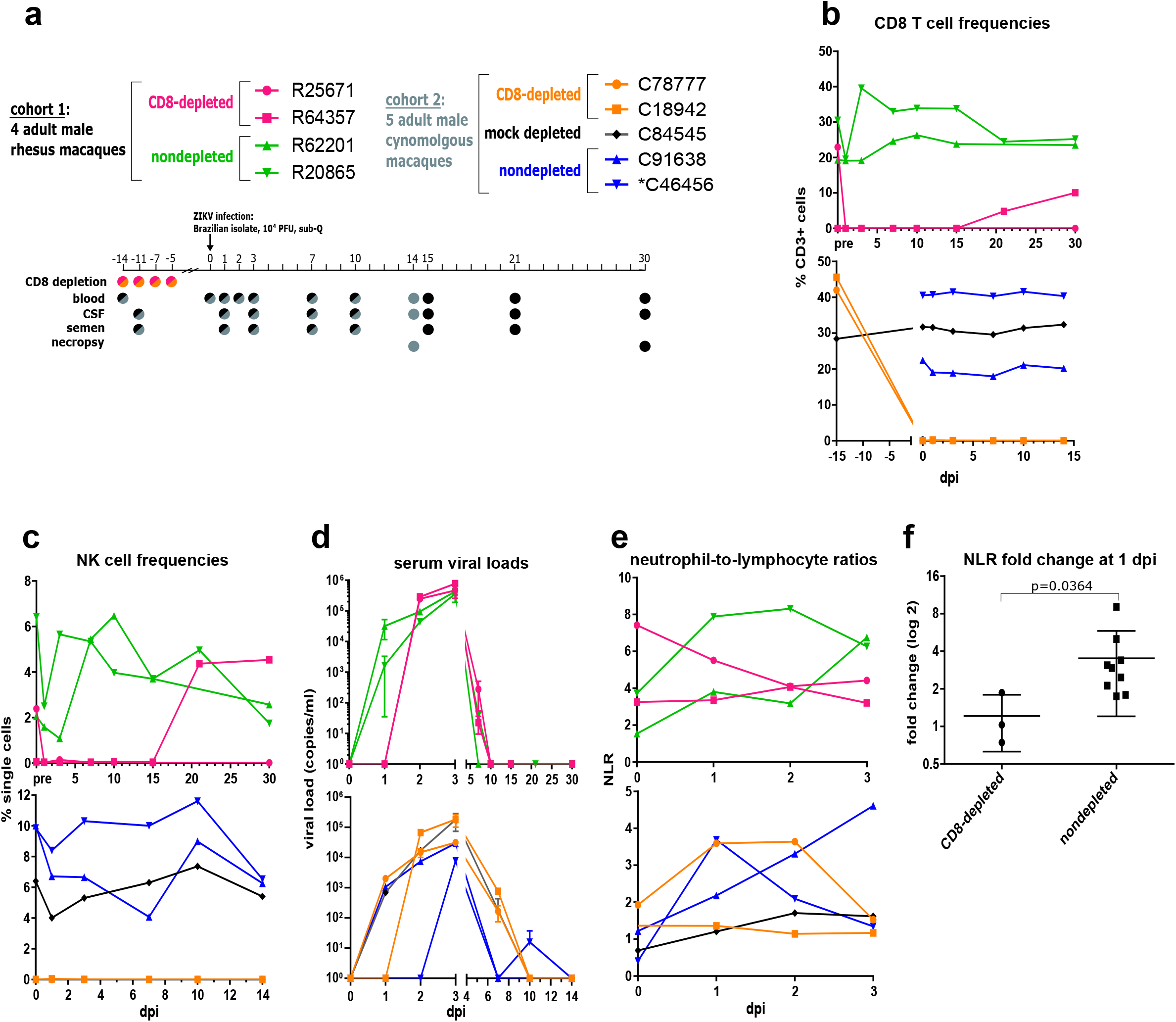
Delayed serum viremia and altered leukocyte kinetics in CD8-depleted macaques. (**A**) Study design. Two animals of each cohort were depleted of CD8+ lymphocytes, and all animals were infected with a Brazilian isolate of ZIKV. Viremia was tracked over 30 days in cohort 1 and 14 days in cohort 2 before necropsy. Colors used in the timeline correspond to cohorts and treatment conditions. C46456 (nondepleted) was splenectomized in a previous study and is indicated by an asterisk (*) throughout. (**B**) Flow cytometric analysis of CD8 T cell frequencies in PBMCs over time. (*Top*): cohort 1; (*Bottom*): cohort 2 (consistent throughout). (**C**) NK cell frequency, as measured by flow cytometry. (**D**) Viral RNA in serum over infection (error bars, SD). (**E**) NLR, derived using total neutrophil and lymphocyte counts in blood from CBC data. (**F**) Fold change in NLR from 0 to 1 dpi among 4 rhesus and 4 cynomolgus macaques included in the present study in addition to a previous cohort of 4 ZIKV-infected non-pregnant female rhesus macaques (center line, mean; error bars, SD). (The blood of one CD8-depleted cynomolgus macaque, C18942, clotted prior to complete blood count (CBC) analysis, precluding calculation of NLR for this animal.) The significant (p ≤ 0.05) difference in NLR fold change was determined using a Mann-Whitney test.

Whole blood, cerebrospinal fluid (CSF), and semen were obtained from animals at the indicated timepoints (Fig. 1a). Peripheral blood mononuclear cells (PBMCs) were isolated from the blood of rhesus macaques using SepMate tubes (Stemcell Technologies) according to the manufacturer’s protocol or from the blood of cynomolgus macaques using Lymphoprep (Stemcell Technologies) for standard density gradient centrifugation. At necropsy, the indicated tissues were collected and snap-frozen.

### Virus quantification

Viral RNA was extracted from serum and CSF using the High Pure Viral RNA Kit (Roche). Semen, as well as the indicated lymphoid, reproductive, GI, and neural tissues were homogenized in Qiazol (Qiagen) using either disposable tissue grinders (Fisherbrand) or a TissueRuptor (Qiagen), and RNA was isolated using the RNeasy Lipid Tissue Mini Kit (Qiagen). Viral RNA from body fluids and tissues was quantified using qRT-PCR as described previously (Magnani et al., 2018b).

### Antiviral gene expression assays

2.5 ml whole blood was drawn from each animal at 0, 1, 3, and 15 dpi into PAXgene blood RNA tubes (PreAnalytiX) and equilibrated to −80°C as per the manufacturer’s protocol. RNA was extracted from blood samples using the PAXgene blood RNA kit (PreAnalytiX), and cDNA was synthesized using the RT2 First Strand Kit (Qiagen). Transcriptional profiles of immune signaling were generated using the nCounter NHP Immunology Panel of 770 macaque immune response genes (NanoString Technologies). In whole blood, transcriptional responses were assessed at 3 dpi relative to expression levels pre-infection using nSolver software v4.0 (NanoString Technologies). Fold change data were imported into Ingenuity Pathway Analysis (IPA) (Qiagen) to discern relevant signaling pathways and disease functions. Antiviral transcriptional responses were confirmed by way of qRT-PCR using a rhesus macaque RT2 Profiler PCR Array (Qiagen). Responses within each species and treatment group were analyzed together to identify expression levels at the indicated timepoints relative to pre-infection. Heatmaps of gene expression and disease-related pathways were generated using Morpheus (https://software.broadinstitute.org/Morpheus). Principal component analysis (PCA) of antiviral gene expression was performed using ClustVis (Metsalu and Vilo, 2015).

To identify cell populations contributing to antiviral signaling in blood, the CD14 and CD8 MicroBead kits (Miltenyi Biotec) were used to sort CD14+ monocytes and CD8+ lymphocytes from the PBMCs of the indicated animals at peak transcriptional activity (3 dpi). RNA was isolated from cell fractions using the RNeasy Mini Kit (Qiagen), and cDNA was synthesized using the RT2 First Strand Kit (Qiagen). Transcriptional profiling was performed using the nCounter NHP Immunology Panel (NanoString) and verified by RT2 qPCR Primer Assays (Qiagen) for ISG15, OAS2, and DDX58.

To characterize antiviral signaling in myeloid cells, PBMCs were isolated from the whole blood of ZIKV-naïve colony rhesus macaques as described above, and the CD14 MicroBead kit (Miltenyi Biotec) was used to isolate monocytes. For the co-culture assay, monocytes were incubated in the presence or absence of autologous CD8+ cells and/or ZIKV in a 96-well U-bottom plate with 50,000 cells of each type plated per well for the indicated timepoints. qPCR for ISG15 and OAS2 was performed as described above. For antiviral gene screening in monocyte-derived macrophages (MDMs), monocytes were cultured at 1 × 10^6^ cells/ml in RPMI-1640 medium supplemented with 1% human AB serum (Sigma), 20 ng/ml M-CSF (Peprotech), 1% L-glutamine, and 1% penicillin/streptomycin. After 7 days of culture, monocytes were sufficiently differentiated into MDMs and were either infected with the same Brazilian ZIKV isolate described above or left uninfected. At 24 hpi, RNA was extracted using the RNeasy Mini Kit (Qiagen), cDNA was synthesized using the RT2 First Strand Kit (Qiagen), and transcriptional signaling was assessed using the rhesus macaque antiviral response RT2 Profiler PCR Array (Qiagen). Antiviral gene expression in ZIKV-infected monocyte-derived macrophages (MDMs) was calculated relative to uninfected controls.

### Flow cytometry and gating strategy

For absolute lymphocyte counts, whole blood was stained within 2 hours of blood draw for the surface markers CD45 (PerCP; DO58-1283; BD Biosciences), CD3 (FITC; SP34; BD Biosciences), CD4 (APC; L200; BD Biosciences), and CD8 (V500; SK1; BD Biosciences). Flow cytometry was performed on a BD FACSVerse instrument, and absolute counts were calculated using FACS Suite software.

For immunophenotyping, PBMCs from the indicated timepoints were thawed, washed, and stained using Live/Dead Fixable Aqua Dead Cell Stain Kit (Invitrogen). PBMCs were then stained for the surface markers CD16 (AL488; 3G8; BioLegend), CD169 (PE; 7-239; BioLegend), CD28 (PECF594; CD28.2; BD Biosciences), CD95 (PCP-Cy5.5; DX2; BioLegend), CD3 (PE-Cy7; SP34-2; BD Biosciences), CD8 (PacBlue; SK1; BioLegend), CD14 (BV605; M5E2; BD Biosciences), HLA-DR (BV650; L243; BioLegend), NKG2A (APC; Z199; Beckman Coulter), and CD4 (APC-H7; L200; BD Biosciences). Cells were subsequently fixed in FluoroFix buffer (BioLegend), permeabilized using Perm/Wash buffer (BioLegend), and stained intracellularly for CD69 (BV711; FN50; BD Biosciences) and Ki67 (AL700; B56; BD Biosciences). Flow cytometry was performed on a BD LSRII instrument and data were analyzed using FlowJo (vX.10.4.2) and visual t-distributed stochastic neighbor embedding (viSNE) (Cytobank) softwares. For viSNE analysis, live singlet monocytes (CD14+ and/or CD16+) or live singlet CD3+ T cells were gated prior to downsampling at a minimum of 500 cells per animal in FlowJo v. 10.5.3. Downsampled files for each animal were then concatenated by group (i.e., species, dpi, and treatment condition). When the number of animals differed per group, concatenated files were further downsampled to achieve an equal number of cells per group. viSNE was conducted using Cytobank with the following settings: Perplexity = 30, Iterations = 1000, Theta = 0.5, Seed =random, Compensation = internal file. For the monocyte viSNE, the following parameters were utilized in the run: Ki67, CD14, HLA-DR, CD69, CD95, CD14, and CD169. For the CD3+ T cells viSNE, the following parameters were utilized in the run: Ki67, CD4, HLA-DR, CD69, CD95, CD28, CD3, CD8.

For general immunophenotyping analysis, cytometry data were first gated for lymphocytes, singlets, and live cells. NK cells were considered as CD8+/CD16+. CD4 T cells (CD3+/CD4+) and CD8 T cells (CD3+/CD8+) were gated into naïve (CD28+/CD95-), central memory (CM) (CD28+/CD95+), and effector memory (EM) (CD28-/CD95+) subsets. CD3-cells were divided into B cells (DR+/CD14-/CD16-) and monocytes (classical, CD14++/CD16-; intermediate, CD14+/CD16+; nonclassical, CD14^low^/CD16+). Cell subsets were analyzed with respect to frequency, proliferation (Ki67+) and activation (CD69+ or CD169+).

### Intracellular cytokine staining

PBMCs from the indicated timepoints were thawed and rested overnight prior to stimulation with peptide pools comprising ZIKV capsid (C), membrane (M), envelope (E), and nonstructural protein 1 (NS1) (BEI Resources). On peptide stimulation, cells were also treated with brefeldin A (BioLegend), GolgiStop (BD Biosciences), anti-CD28 (NHP Reagent Reference Program, www.nhpreagents.org/), anti-CD49d (9F10; BioLegend), and anti-CD107a (AL700; H4A3; BD Biosciences). 24 hours post-stimulation, cells were stained for the surface markers CD3 (PE-Cy7; SP34-2; BD Biosciences), CD8 (PacBlue; SK1; BioLegend), and CD4 (APC-H7; L200; BD Biosciences). Cells were also fixed and permeabilized as described above and stained intracellularly for perforin (FITC; Pf-344; Mabtech), granzyme B (PE; GB12; Invitrogen), CD69 (PE-CF594; FN50; BD Biosciences), IL-2 (PCP-Cy5.5; MQ1-17H12; BD Biosciences), and IFNg (AL647; 4S.B3; BioLegend). Flow cytometry was performed on a BD LSRII instrument and data were analyzed using FlowJo software (vX.10.4.2).

### Plaque reduction neutralization tests

ZIKV plaque reduction neutralization tests (PRNTs) were conducted according to previously published protocols (Lieberman et al., 2009; Ward et al., 2018). Briefly, ZIKV MEX-I-44 isolated in Tapachula, Mexico in 2016 was obtained from The University of Texas Medical Branch, Galveston, TX and cultured to passage 8 in Vero cells. Serum specimens were incubated for one hour at serial dilutions of 1:10, 1:20…1:320 with a previously frozen virus stock of known plaque forming unit (PFU). Samples were then inoculated in duplicate onto a mono-layer of Vero cells grown on 6-well plates and allowed to incubate for an additional hour. Infectious material was then removed and replaced with a 1:1 mixture of Vero media and Avicel^®^ before being incubated for 4 days. To read plaques, the Avicel^®^ layer was fixed with 10% neutral buffered formalin. Finally, the formalin-Avicel^®^ layer was removed and the monolayer was stained with crystal violet, washed with tap water and allowed to dry before plaques were counted manually.

Percent reduction in observed plaques and a PRNT90 cutoff were used for interpretation. A PRNT90 titer is the dilution of a sample at which a 90% reduction in possible plaques is observed. The maximum number of potential plaques was obtained for each run using a corresponding back-titration and a linear model was fit to the observed number of plaques for each dilution. A PRNT90 titer was derived for each sample using the linear model and the equation for a straight line in the statistical program R (R Core Team, 2018). For samples that were positive but above the resolution of the PRNT assay the value of the greatest number of possible plaques for that run, as determined by the back titration, was assigned for each dilution for use with the linear model.

### Histology

Tissues samples collected at necropsy were fixed in Z-Fix (Anatech), embedded in paraffin and 5 *μ* m thick sections were cut, adhered to charged glass slides, and either stained routinely with hematoxylin and eosin or Prussian blue.

### Statistical analysis

Statistical analysis was conducted using GraphPad Prism v6.07 (GraphPad Software). A Mann-Whitney test was used to compare neutrophil-to-lymphocyte ratios (NLRs) among treatment conditions (CD8-depleted versus nondepleted).

## Results

### Delayed serum viremia and altered leukocyte kinetics

CD8+ lymphocyte depletion commenced 14 days prior to ZIKV infection (Fig. 1a), and CD8 T cells were undetectable in all depleted animals well before infection (Fig. 1b & Supplementary Fig. 1a). To achieve CD8 depletion, we used the MT807R1 antibody (Schmitz et al., 1999) to target CD8α, effectively depleting CD8 T cells (Fig. 1b & Supplementary Fig. 1a), CD8+/CD4+ double-positive T cells (Supplementary Fig. 1b), and NK cells (Fig. 1c) but not CD4 T cells (Supplementary Fig. 1c) from the blood of all treated animals. As MT807R1 targets NK cells in addition to CD8+ lymphocytes, any deficiency in host response following treatment with anti-CD8α could indicate that either or both types of cells are important for acute control of ZIKV. Flow cytometric analysis of NK cell frequency in nondepleted animals revealed expansion early in infection (Fig. 1c). Intriguingly, the CD8-depleted macaque R64357 recovered CD8 T cells and NK cells at later timepoints, between 15 and 21 days post-infection (dpi) (Figs. 1b-c).

Following subcutaneous inoculation with ZIKV, nondepleted animals and a single CD8-depleted cynomolgus macaque experienced rapid serum viremia of 3-4.5 logs at 1 dpi (Fig. 1d), consistent with previous reports of ZIKV in both rhesus and cynomolgus macaques (Dudley et al., 2016; Koide et al., 2016; Osuna et al., 2016b). Strikingly, serum viremia in 3 of 4 CD8-depleted macaques was delayed until 2 dpi, when viral RNA was higher than in nondepleted animals (Fig. 1d). Viremia was also delayed until 3 dpi in *C46456 (Fig. 1d), a nondepleted cynomolgus macaque that had been previously splenectomized. Perhaps importantly, the spleen is a major site of replication and spread of the related mosquito-borne flaviviruses dengue virus (DENV) (Prestwood et al., 2012) and West Nile virus (WNV) (Bryan et al., 2018) and is also an immense reservoir of monocytes (Swirski et al., 2009), which are permissive to ZIKV replication in humans (Michlmayr et al., 2017) and macaques (O’Connor et al., 2018). The lack of a spleen in *C46456 (nondepleted) might have precluded ZIKV replication in this important target organ, thereby delaying viral kinetics.

By 7 dpi, serum viremia persisted in the CD8-depleted cynomolgus macaques in addition to the mock-depleted control, while viremia was undetected in both nondepleted animals of the cohort (Fig. 1d). For the remainder of the study, viral kinetics were similar among cohorts and treatment conditions, peaking at 3 dpi and dropping to undetectable levels by 10 dpi and beyond (Fig. 1d). This was again in exception to *C46456 (nondepleted), which showed a small viral rebound at 10 dpi. A previous cohort of female macaques infected with the identical strain of ZIKV demonstrated similar patterns of serum viremia to those observed in nondepleted animals (Supplementary Fig. 2a).

The previous female cohort also showed consistent patterns of innate immune cell recruitment in the blood one day following ZIKV infection, summarized by the biomarker neutrophil-to-lymphocyte ratio (NLR) (Faria et al., 2016). Patterns of leukocyte mobilization included a spike in neutrophil frequency (Supplementary Fig. 2c) and a simultaneous drop in lymphocyte frequency (Supplementary Fig. 2d), resulting in an elevated NLR at 1 dpi (Supplementary Fig. 2e). These findings were generally recapitulated in nondepleted animals but not in CD8-depleted animals (Figs. 1e-f and Supplementary Figs. 2c-e), potentially linking NLR and acute serum viremia, although the trend did not hold in the mock-depleted cynomolgus macaque C84545. This association is strengthened by prototypical patterns of leukocyte homing in C78777 (Fig. 1e), the only CD8-depleted macaque with serum viremia at 1 dpi (Fig. 1d).

### Differential monocyte-driven transcriptional profiles

To characterize immune responses that might be differentiating patterns of viremia and leukocyte mobilization, we used the NanoString platform to quantify the expression of macaque immune-related genes in whole blood. CD8-depleted and nondepleted animals showed highly divergent profiles in genes related to IFNα signaling (Fig. 2a) and leukocyte homing (Supplementary Fig. 3a), resulting in the robust induction of disease-related pathways in nondepleted macaques that failed to activate in animals lacking CD8+ lymphocytes (Supplementary Fig. 3b). To confirm the transcriptional quiescence evident in CD8-depleted macaques, we used a quantitative real-time PCR (qRT-PCR) array of 84 antiviral genes in the rhesus macaque genome. Consistent with the NanoString results, nondepleted rhesus and cynomolgus macaques showed strong induction of several RIG-I like receptors (RLRs) and type-I IFN stimulated genes (ISGs) at 3 dpi, synchronous with peak serum viremia (Fig. 2b). The most highly induced genes include the pattern recognition receptors *TLR3, DDX58* (also known as *RIG-I*), and *IFIH1* (also known as *MDA5*), as well as the ISGs *ISG15, MX1*, and *OAS2*. Principal component analysis (PCA) of antiviral signaling further discriminated the transcriptional phenotypes in CD8-depleted and nondepleted animals (Fig. 2c). The induction of antiviral genes in nondepleted rhesus macaques was highest at 3 dpi and was generally followed by a return to near-baseline expression by 15 dpi (Supplementary Fig. 3c). In contrast to nondepleted animals, CD8-depleted macaques showed a virtual absence of transcriptional responses in whole blood at all timepoints tested (Figs. 2a-b and Supplementary Fig. 3d).

**Figure 2.**
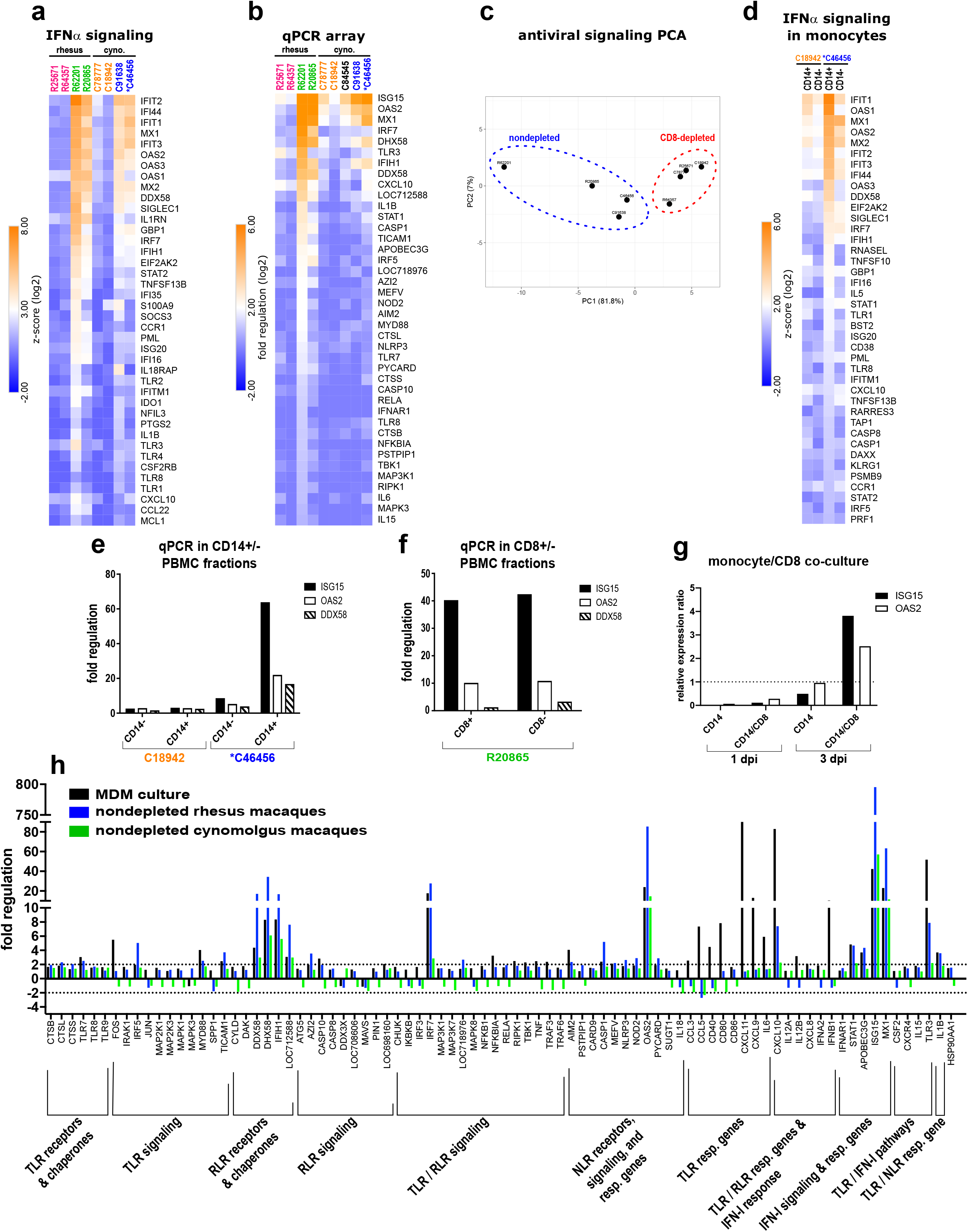
Differential monocyte-driven transcriptional profiles among CD8-depleted and nondepleted macaques. (**A**) Activation of downstream IFNα effector molecules in whole blood at 3 dpi relative to pre-infection (cyno. = cynomolgus, consistent throughout). **(B)** Fold regulation of antiviral gene expression in whole blood at 3 dpi, confirmed using a qRT-PCR array of 84 genes in the rhesus macaque genome (dep. = CD8-depleted; non. = nondepleted). **(C)** PCA of antiviral gene expression in whole blood at 3 dpi. **(D)** Activation of downstream IFNα effector molecules in sorted CD14+ monocytes and CD14-PBMCs at 3 dpi. **(E)** qPCR confirmation of antiviral gene expression in CD14+ monocytes and CD14-PBMCs at 3 dpi. Gene induction is normalized to b-actin, and fold regulation is expressed relative to 0 dpi. **(F)** Antiviral gene expression in CD8+ and CD8-fractions of PBMCs from a nondepleted rhesus macaque at 3 dpi. Gene induction is normalized to b-actin, and fold regulation is expressed relative to 15 dpi. **(G)** Antiviral gene expression in co-cultured CD14+ monocytes and autologous CD8+ cells from ZIKV-naïve PBMC infected with ZIKV *ex vivo* at 1 and 3 dpi. **(H)** Comparison of antiviral gene induction in cultured, ZIKV-infected MDMs (red) and the whole blood of nondepleted rhesus (blue) and cynomolgus (green) macaques at 3 dpi. Genes included in the qPCR array relate to toll-like receptor (TLR), nod-like receptor (NLR) or type-I interferon (IFN-I) signaling (resp. = responsive).

Although we suspected monocytes to be driving antiviral gene expression owing to their susceptibility to ZIKV infection (Michlmayr et al., 2017; O’Connor et al., 2018), an important caveat of probing whole blood is that the identity of the cell populations responding transcriptionally is unknown. To resolve cell populations contributing to antiviral signaling in blood, we sorted CD14+ monocytes from peripheral blood mononuclear cells (PBMCs) at 3 dpi and profiled their expression of immune genes. Probe hybridization revealed selective antiviral gene expression in the monocytes of a nondepleted macaque and showed that even purified and sorted PBMCs from a CD8-depleted animal fail to establish transcriptional responses at peak serum viremia (Fig. 2d). These findings were verified by qRT-PCR (Fig. 2e). Nonetheless, it remained possible that the lack of a transcriptional response in CD8-depleted macaques could have been attributed in part to an absence of otherwise responding NK cells. Sorting PBMCs from a nondepleted rhesus macaque into CD8+ and CD8-fractions, we found similar levels of gene induction in both populations, although expression was marginally higher in the CD8-subset, and transcription of *DDX58* was almost exclusive to CD8-cells (Fig. 2f). Gene induction in CD8+ cells indicates that NK cells may indeed contribute to antiviral signaling, but similar transcriptional activation in the CD8-fraction affirms that the absence of transcriptional activation in CD8-depleted animals was not simply the product of a lack of NK cells.

To further explore transcriptional responses to ZIKV in myeloid cells, we infected monocyte-CD8 cell co-cultures *ex vivo* and compared antiviral gene expression to monocytes infected in the absence of CD8+ cells, finding that the presence of CD8+ lymphocytes is important in promoting transcriptional responses (Fig. 2g). We also cultured monocyte-derived macrophages (MDMs) *in vitro*, infected the macrophages with ZIKV, and profiled antiviral gene expression using qRT-PCR. We found an overlapping transcriptional fingerprint to those observed in the blood of nondepleted rhesus and cynomolgus macaques at 3 dpi (Fig. 2h), suggesting that ZIKV-permissive myeloid cells may be driving antiviral gene induction *in vivo*. Although cultured MDMs exhibited higher induction of several TLR responsive genes (Fig. 2g), this difference might be attributed to cell type.

### Altered monocyte activation and frequency

Divergent transcriptional patterns in CD8-depleted and nondepleted macaques could be induced by differentially responding monocytes, given that monocytes are known targets of ZIKV infection (Lum et al., 2018; O’Connor et al., 2018) and contribute to antiviral signaling during ZIKV infection (Lum et al., 2018). To interrogate the immunophenotypic effects of CD8 depletion, we developed a multicolor flow cytometry panel to track innate and adaptive immune cells over time. The resulting data were highly dimensional, comprising a variety of surface markers and sampling animals at multiple timepoints and with respect to different treatment groups. To survey general immune responses over time, we used an adaptation of t-distributed stochastic neighbor embedding (tSNE), viSNE (Amir et al., 2013).

In both rhesus and cynomolgus macaques, CD8 depletion dysregulated the kinetics of monocyte activation as measured by CD169 (siglec-1) expression (Biesen et al., 2008; Hirsch et al., 2018; York et al., 2007). Nondepleted rhesus and cynomolgus macaques showed early activation of monocytes, peaking at 3 dpi and returning sharply to baseline by 14-15 dpi (Figs. 3a-c). Upregulation of CD169 in nondepleted animals was affirmed at the RNA level (Fig. 2a). Although patterns of CD169 induction were consistent in all monocyte subsets (Supplementary Figs. 4a-c), viSNE analysis indicated that CD169 was most highly expressed on intermediate and nonclassical monocytes in both cohorts (Figs. 3a-b). Contrasting nondepleted animals, CD8-depleted rhesus and cynomolgus macaques showed less well-defined monocyte activation at 3 dpi, which was accompanied in rhesus macaques by prolonged monocyte activation beyond 15 dpi (Figs. 3a-c & Supplementary Figs. 4a-d). These findings were consistent transcriptionally, as whole blood from CD8-depleted animals had muted expression of genes related to myeloid cell activation (Fig. 3d). Monocyte subsets showed additional nuances in phenotype that appeared dependent on CD8 depletion: In rhesus macaques, CD95 (Fas) was increased on classical monocytes in CD8-depleted animals (Supplementary Fig. 4e) and on nonclassical monocytes in nondepleted animals (Supplementary Fig. 4f), although similar patterns were not observed in cynomolgus monkeys.

**Figure 3.**
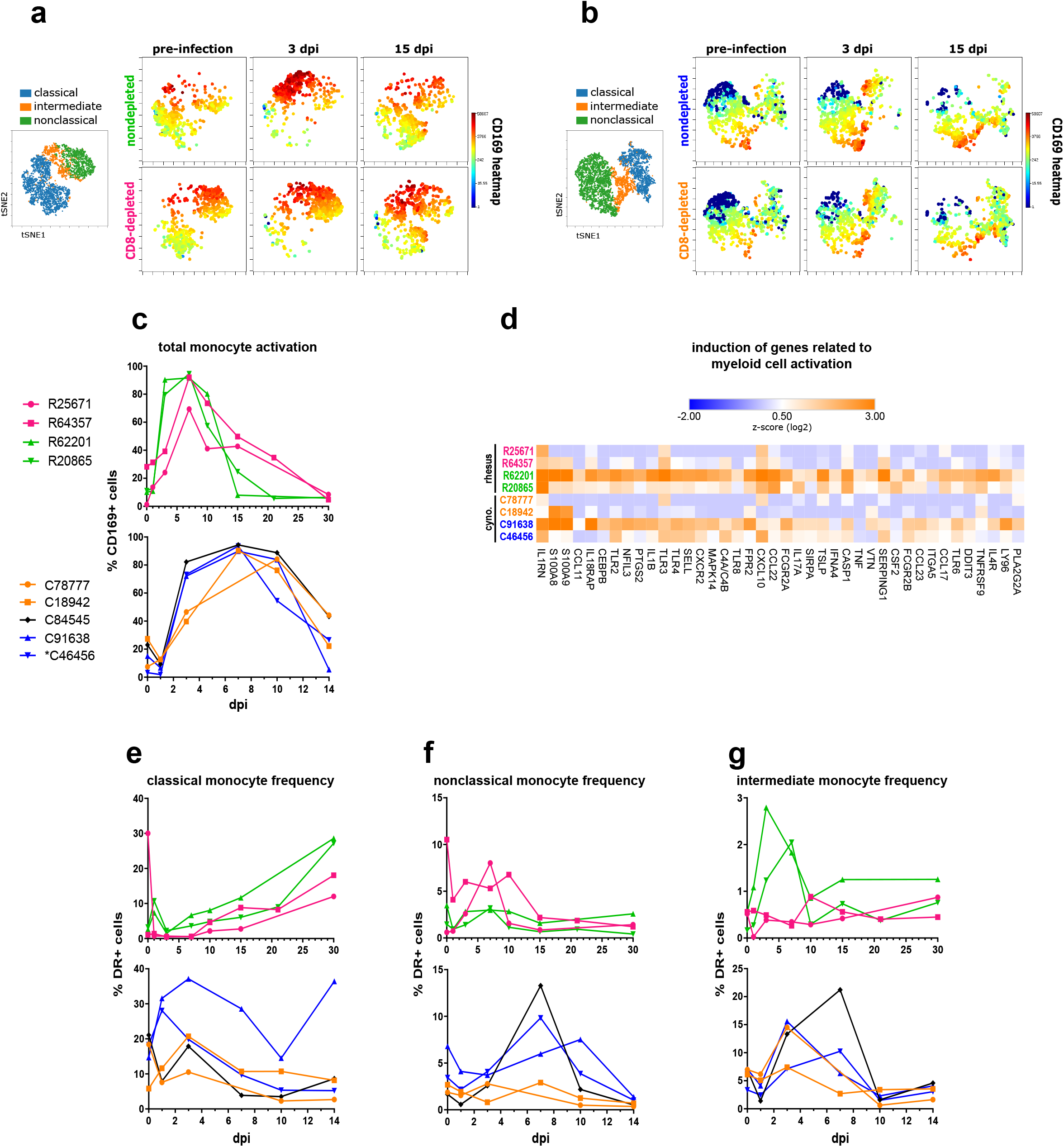
Altered monocyte activation and frequency in CD8-depleted macaques. (**A-B**) viSNE analyses of monocyte activation in rhesus (A) and cynomolgus (B) macaques, as measured by CD169 mean fluorescence intensity (MFI). Dot plots are concatenated for animals within each treatment condition. The viSNE clustering profile of monocyte subsets (*left*) correspond to cell populations in the CD169 MFI heatmaps in nondepleted and CD8-depleted animals over time (*right*). **(C)** summaries of CD169 expression in total monocytes. **(D)** Induction of genes related to myeloid cell activation at 3 dpi relative to pre-infection. (**E-G**) Frequencies of classical (E), nonclassical (F), and intermediate (G) monocyte subsets in rhesus and cynomolgus macaques over time.

CD8 depletion also differentially modulated the abundance of monocyte subsets in blood. One day following ZIKV infection, classical monocytes expanded immediately in nondepleted animals of both cohorts (Fig. 3e) excluding the mock-depleted cynomolgus macaque C84545. During acute infection (3-7 dpi), the frequency of nonclassical monocytes increased preferentially in CD8-depleted rhesus macaques and in nondepleted cynomolgus macaques (Fig. 3f). Nondepleted rhesus macaques showed an expansion of intermediate monocyte frequency at 3-7 dpi (Fig. 3g), although a CD8-dependent effect on intermediate monocyte frequency was not evident in cynomolgus monkeys.

### Compensatory adaptive immune responses

Adaptive immune responses to ZIKV were also differentially modulated by CD8 depletion, with apparent compensatory responses in CD8-depleted macaques of both cohorts. Nondepleted rhesus and cynomolgus macaques began to mount CD8 T cell responses 7-10 dpi, which were characterized by proliferation (Ki67) and activation (CD69) of effector memory (EM), central memory (CM), and naïve CD8 T cell subsets (Figs. 4a-c & Supplementary Figs 5a-e). These responses were antigen-specific and functional, given that CD8 T cells stimulated with ZIKV peptides produced IFNg and contained perforin by intracellular cytokine staining (ICS) (Fig. 4e). Intriguingly, the CD8-depleted rhesus macaque R64357 also showed evidence of a CD8 T cell response at 21 dpi (Figs. 4c & 4e), concomitant with the recovery of CD8+ lymphocytes in this animal (Supplementary Fig. 1a). Although tested, antigen-specific T cell responses were not detected by ICS in cynomolgus macaques.

**Figure 4.**
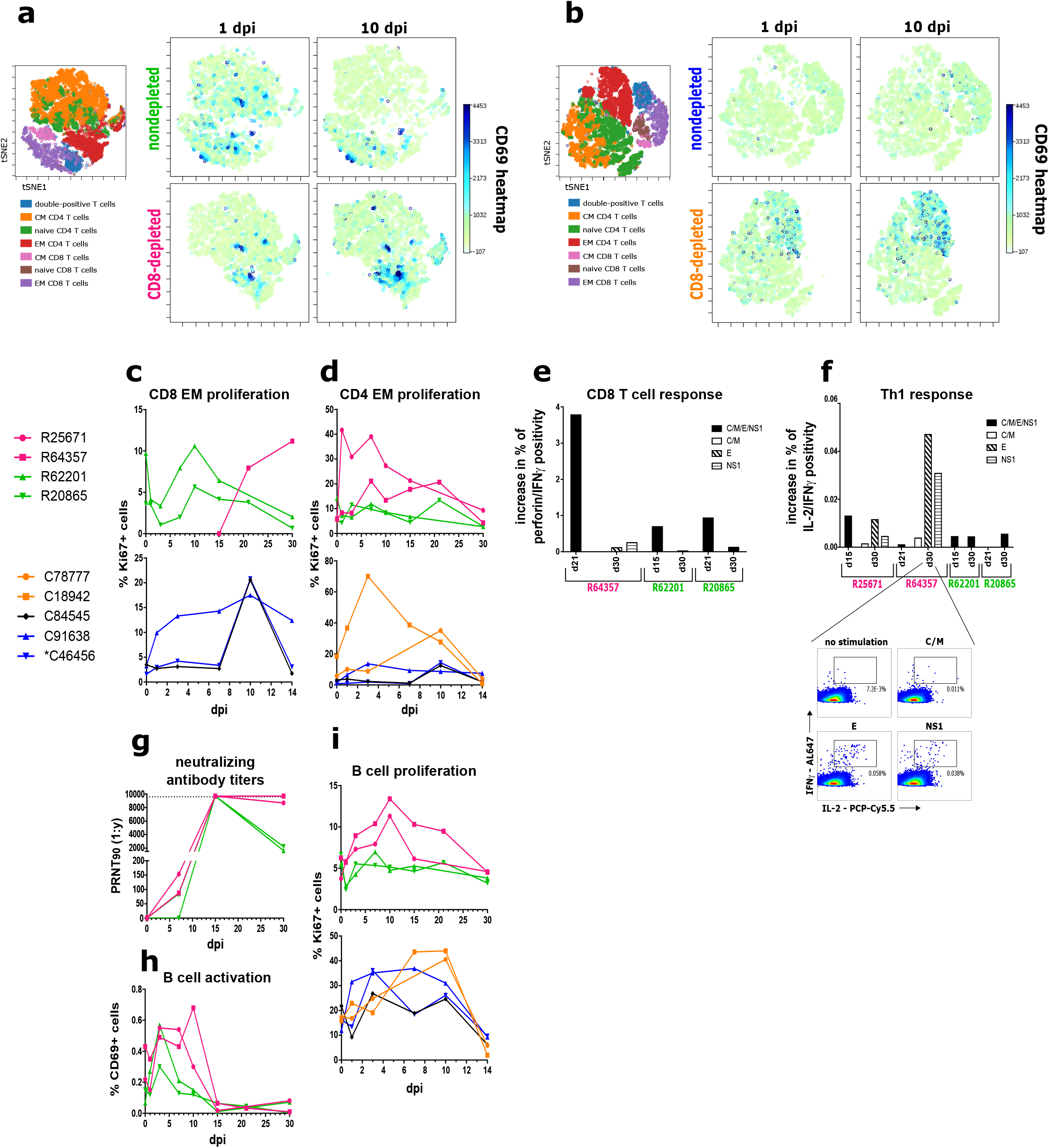
Compensatory adaptive immune responses in CD8-depleted macaques. (**A-B**) viSNE analyses of T cell activation in rhesus (A) and cynomolgus (B) macaques, as measured by CD69 expression. Dot plots are concatenated for animals within each treatment condition. The viSNE clustering profiles of CD4 and CD8 T cell subsets (*left*) correspond to cell populations in the CD69 heatmaps in nondepleted and CD8-depleted animals at 1 & 10 dpi (*right*). (**C**) Proliferation of EM CD8 T cells in rhesus and cynomolgus macaques over time. CD8-depleted macaques are excluded owing to the absence of CD8+ lymphocytes in these animals exclusive of R64357 at later timepoints. (**D**) Proliferation of EM CD4 T cells in rhesus and cynomolgus macaques over time. **(E)** CD8 T cell responses in rhesus macaques, assessed by ICS of PBMCs stimulated with viral peptides derived from the indicated ZIKV proteins (C = capsid; M = membrane; E = envelope; NS1 = nonstructural protein 1, consistent throughout). CD8 T cell responses were identified by co-positivity for perforin and IFNγ. (**F**) Th1 responses, determined by ICS for IL-2 and IFNγ co-positivity. (*Inset*): representative antigen-specific cytometry plots for R64357 (CD8-depleted) at 30 dpi. (**G**) Serum neutralizing antibody titers in rhesus macaques, represented as PRNT90. **(H)**Activation of B cells in rhesus macaques over time. **(I)** Proliferation of B cells in rhesus and cynomolgus macaques over time.

The absence of CD8 T cells in depleted rhesus and cynomolgus macaques appeared to promote the reciprocal activation and expansion of CD4 T cell subsets (Figs. 4a-b, 4d and Supplementary Figs. 5f-j), mirroring the kinetics of CD8 T cell activation in nondepleted animals. CD8-depleted rhesus macaques began to induce CD4 T cell responses consistent with a Th1 phenotype, characterized by co-positivity for IL-2 and IFNg, but such responses were not present in nondepleted animals (Fig. 4f).

To gauge humoral responses to ZIKV, we conducted plaque reduction neutralization tests (PRNTs) using rhesus macaque sera to quantify neutralizing antibody titers. All animals except R20865 (nondepleted) showed evidence of neutralizing antibodies at 7 dpi, the earliest post-infection timepoint tested (Fig. 4g). While highly neutralizing titers were detected in all animals at 15 dpi, antibody concentrations declined in nondepleted animals, but not in CD8-depleted animals, beyond this timepoint. Strikingly, depleted rhesus macaques retained highly neutralizing antibody titers until necropsy, a finding consistent with elevated B cell activation (Fig. 4h) and proliferation (Fig. 4i) in these animals.

### Enhanced tissue dissemination and neuropathology

Given the persistence of high neutralizing antibody titers in CD8-depleted rhesus macaques, we suspected that virus might be lingering in the peripheral tissues of these animals. The duration of infection before necropsy differed among the rhesus (30 dpi) and cynomolgus (14 dpi) macaque cohorts to identify patterns of viral dissemination and clearance over time. Informed by previous reports of ZIKV tropism in macaques (Coffey et al., 2017; Hirsch et al., 2017; Osuna et al., 2016b), we searched for viral RNA in lymphoid, neural, gastrointestinal (GI), and reproductive tissues, as well as in semen and cerebrospinal fluid (CSF) to evaluate viral distribution in these sites.

Relative to nondepleted animals, CD8-depleted cynomolgus macaques had markedly higher levels of ZIKV RNA in the inguinal, mesenteric, and colonic lymph nodes, as well as in the spleen and jejunum (Figs. 5a and 5c). All cynomolgus monkeys except C91638 (nondepleted) harbored virus in the rectum without an obvious difference among treatment groups. Notably, the trend of higher viral burdens in the lymphatic tissues of CD8-depleted animals was consistent in rhesus macaques (Fig. 5f). CD8 depletion also appeared to promote ZIKV dissemination in the semen, with both CD8-depleted cynomolgus macaques presenting semen viral loads and no viral RNA detected in nondepleted animals of the same cohort (Fig. 5d). Intriguingly, the nondepleted macaque C84545 (mock-depleted) showed the highest level of viral RNA in the prostate and was the only animal to present virus in the testes (Fig. 5c), yet no ZIKV was detected in the semen of this animal (Fig. 5d). These findings too were consistent in rhesus macaques, with virus detected in the semen (Fig. 5g) and seminal vesicle (Fig. 5f) of a CD8-depleted animal and only a miniscule quantity of viral RNA detected in the semen of a nondepleted animal (Fig. 5g).

**Figure 5.**
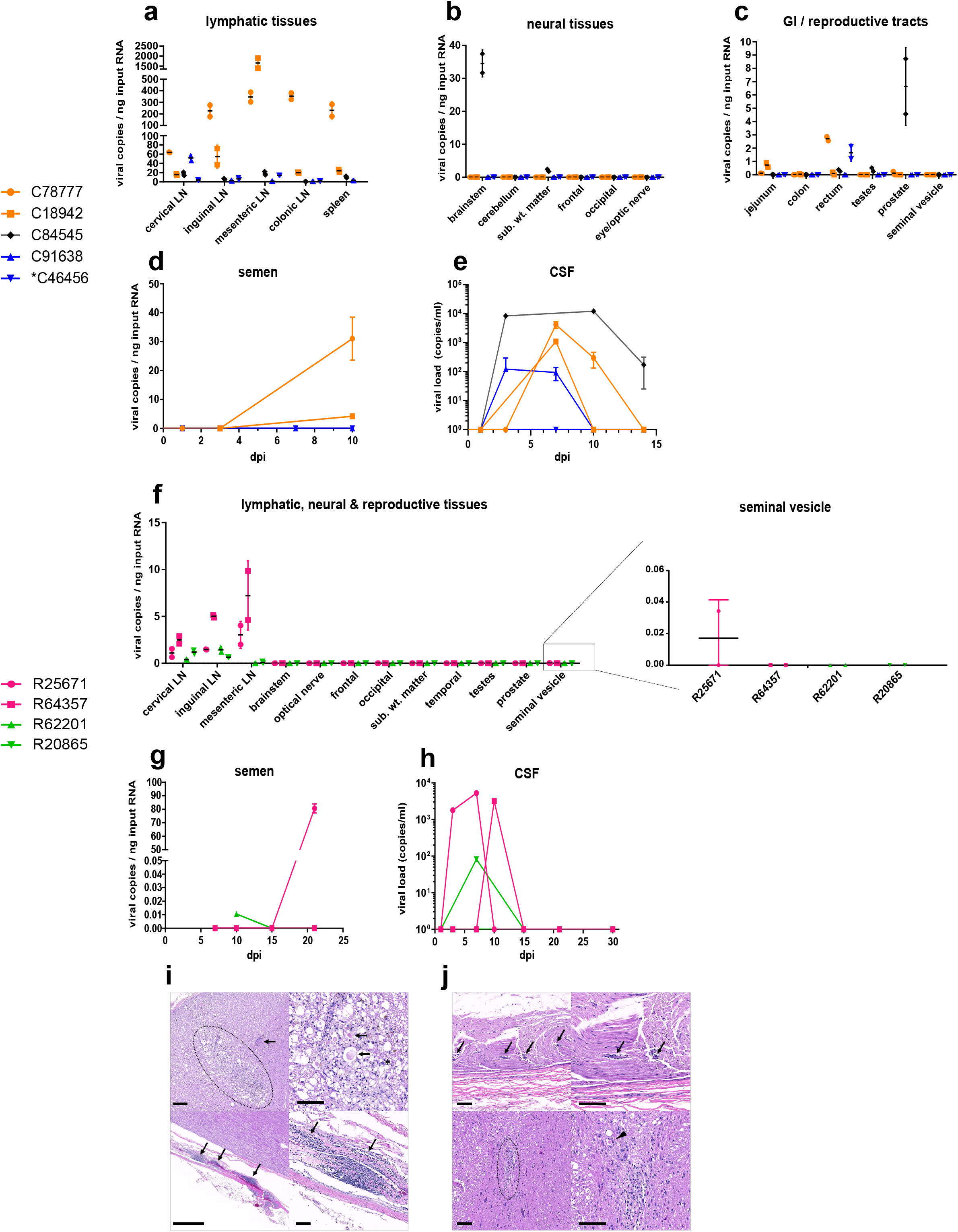
Enhanced tissue dissemination and neuropathology in CD8-depleted macaques. (**A-E**) Viral dissemination in cynomolgus macaques, including lymphatic (A), neural (B) and reproductive (C) tissues, as well as semen (D) and CSF (E) (center line, mean; error bars, SD of two replicates per sample, LN = lymph node; sub. wt. matter = subcortical white matter, consistent throughout). **(F-H)** Viral dissemination in rhesus macaques, including lymphatic, neural, and reproductive tissues (F), as well as semen (G) and CSF (H). (**I**) R25671 (CD8-depleted) brainstem (top) and lumbar spinal cord (bottom). *Top:* There is an area of encephalomalacia (dotted region, left) adjacent to a vessel that exhibits medial thickening (arrow, left). The area of malacia is characterized by dilated myelin sheaths with swollen axons (arrow, right) or gitter cell infiltration (asterisks, right). H&E, Bar = 100 um. *Bottom:* The meninges surrounding the lumbar spinal cord are multifocally infiltrated by aggregates of lymphocytes (arrows). H&E, Bar = 1 mm (left) and 100 um (right). (**J**) R64357 (CD8-depleted) sciatic nerve (top) and brainstem lesions (bottom). *Top*: Small vessels within the sciatic nerve are surrounded by low numbers of lymphocytes (arrows). *Bottom:* A focal glial nodule is present within the gray matter of the brainstem (dotted region, left) with dilation of adjacent myelin sheaths and spheroid formation (arrowhead, right). H&E, Bar = 100 um.

ZIKV RNA was detected in the brainstem and subcortical white matter of C84545 (mock-depleted) (Fig. 5b), and this animal also presented a high magnitude viral load in the CSF early in infection, which persisted until necropsy (Fig. 5e). Exclusive of C84545 (mock-depleted), CD8-depleted cynomolgus and rhesus macaques manifested CSF viral loads at least an order of magnitude greater than nondepleted animals (Figs. 5e and 5h). Although ZIKV was not detected in the central nervous system (CNS) of any rhesus macaque, R25671 (CD8-depleted) and R64357 (CD8-depleted) manifested neural lesions at necropsy that were not present in nondepleted animals. Most strikingly, the brainstem of R25671 showed an area of severe multifocal to coalescing encephalomalacia which showed evidence of Wallerian degeneration, characterized by vacuolation, swollen axons, and infiltration by lymphocytes and phagocytic gitter cells (Fig. 5i). Gitter cells are occasionally found within dilated myelin sheaths. Scant brown granular pigment (presumed hemosiderin) and a proliferative cerebral vessel adjacent to the malacia may indicate that the malacia is the result of a vascular event (thromboembolism, infarct, ischemia, etc.). Additionally, lymphocytic infiltrate was present in the meninges surrounding the lumbar spinal cord (Fig. 5i). No gross abnormalities were noted in R64357, although the sciatic nerve exhibited mild lymphocytic perivasculitis. The sciatic nerve is a known site of ZIKV replication in mice depleted of CD8 cells (Elong Ngono et al., 2017). Further, the brainstem contained a localized area of gliosis, an indicator of CNS damage (Garman, 2011), and dilated myelin sheaths (Fig. 5j). A cause for these neural inflammatory lesions was not apparent by histology.

## Discussion

Owing to the importance of CD8+ T cells in the control of ZIKV in mice (Elong Ngono et al., 2017; Huang et al., 2017), our aim was to explore whether these findings are consistent in nonhuman primates. Despite a small sample size in two cohorts, the absence of CD8+ lymphocytes prompted immediately observable host responses that diverged from previously consistent patterns of viremia and immunity. Contrasting similar experiments with simian immunodeficiency virus (SIV) (Jin et al., 1999; Klatt et al., 2010), the absence of CD8+ lymphocytes did not overtly affect the control of serum viremia. However, the delay of serum viremia in CD8-depleted macaques stood in patent contrast to patterns of acute ZIKV infection observed by our own group (Supplementary Fig. 2a) and others (Dudley et al., 2016). Although a mechanism underlying the delayed serum viremia remains obscure, it is possible that a lack of NK cell stimulation in CD8-depleted animals may be misfiring viral replication in what would otherwise be readily permissive monocytes. Indeed, monocytes engage in intercellular crosstalk with NK cells (Dalbeth et al., 2004; Michel et al., 2012), and IFNg is shown to support ZIKV replication (Chaudhary et al., 2017). NK cell-derived IFNg may activate ZIKV infected myeloid cells in nondepleted animals, promoting an inflammatory milieu that favors early viral replication. Alternatively, NK cells are shown to be minor reservoirs of ZIKV RNA in infected humans (Michlmayr et al., 2017) and pigtail macaques (O’Connor et al., 2018), so the absence of this potential target cell may contribute to the delayed serum viremia in CD8-depleted animals. Additional co-culture assays may elucidate intercellular dynamics important for maintaining patterns of innate immune regulation.

CD8 depletion also impacted the mobilization of leukocyte populations acutely following infection, contrasting patterns reliably observed by our own group (Figs S2c-e) and others (Osuna et al., 2016b). Macaques depleted of CD8 T cells and NK cells show little fluctuation in the biomarker of inflammation NLR, indicating altered innate immune responses immediately following infection. In line with these observations, mice lacking NK cells exhibit altered neutrophil recruitment in a variety of infectious and noninfectious conditions (Costantini and Cassatella, 2011). Neutrophil effector functions are modulated by NK cell-derived cytokines (Costantini and Cassatella, 2011), a signaling axis which might have been disrupted by the depletion of NK cells in macaques.

Confirming miscommunication within the innate immune system of CD8-depleted macaques, these animals presented largely muted transcriptional activity in key virus response pathways during acute infection. In nondepleted macaques, antiviral gene expression was driven principally by circulating monocytes, which stood in sharp contrast to the transcriptional void evident in CD8-depleted animals. Importantly, this signaling pattern was replicated in a co-culture model *ex vivo*. Intercellular crosstalk between monocytes and NK cells affects transcriptional responses to ZIKV infection (Lum et al., 2018), so the absence of NK cells in CD8-depleted macaques might have permitted the infection to evade transcriptional induction in monocytes, complementing the delayed viremia in these animals.

Consistent with a model of monocyte-dependent outcomes in acute ZIKV infection, CD8-depleted and nondepleted macaques also differed in the magnitude and phenotype of their monocyte responses during acute infection. Intermediate and nonclassical monocytes showed the greatest degree of activation, agreeing with recent findings that these subsets are primary targets of ZIKV in the blood (Foo et al., 2017; Jurado and Iwasaki, 2017; Michlmayr et al., 2017). CD8 depletion also impacted the activation of monocytes temporally, further underscoring dysregulated innate responses in depleted animals. CD169 (siglec-1) is a sialic acid-binding lectin previously found to be upregulated in acute ZIKV infection in rhesus macaques (Hirsch et al., 2018; Hirsch et al., 2017). CD169 has important roles in virus capture by myeloid cells (Sewald et al., 2015) and in the mounting of CD8 T cell responses in viral infection (van Dinther et al., 2018), so the robust induction of CD169 in nondepleted animals might have promoted sufficient CD8 T cell responses. CD8 depletion also affected monocyte frequency, possibly contributing to differential transcriptional responses. The transient increase in classical monocytes of both cohorts may be analogous to the monocytosis that accompanies acute ZIKV replication in human patients (Michlmayr et al., 2017). The increase in CD16+ nonclassical monocytes in CD8-depleted rhesus macaques is an outcome also observed in ZIKV infection of human blood (Foo et al., 2017), and the expansion of intermediate monocytes in nondepleted animals resembles ZIKV infection in Nicaraguan patients (Michlmayr et al., 2017). Although minor species differences were evident in the immunophenotyping of rhesus and cynomolgus monocytes, our data support a CD8+ lymphocyte-dependent effect in these transitions, possibly accounting for divergent transcriptional responses in blood.

In addition to modulating innate immune responses, the depletion of CD8+ lymphocytes also promoted compensatory adaptive responses to ZIKV in both cohorts. Nondepleted animals and even the CD8-recovering rhesus macaque R64357 mounted robust CD8 T cell responses, affirming the importance of CD8+ lymphocytes in acute infection. There is precedence for CD8 T cell responses to ZIKV in mice (Elong Ngono et al., 2017; Huang et al., 2017; Pardy et al., 2017) and humans (Grifoni et al., 2018; Grifoni et al., 2017), which appears to be consistent in nonhuman primates. Meanwhile, the presence of Th1 responses and prolonged humoral responses in CD8-depleted animals possibly compensate for the absence of CD8 surveillance. Such adaptive responses are reported in mice, as Th1 polarization (Pardy et al., 2017), and CD4-driven humoral responses (Lucas et al., 2018) are important for controlling infection. Our data support the importance of these adaptive responses in nonhuman primates, especially when the CD8 arm of adaptive immunity is compromised.

The persistence of high neutralizing antibody titers until necropsy in CD8-depleted rhesus macaques also suggested that there might be virus lingering in the peripheral tissues of these animals. Indeed, ZIKV RNA was generally more abundant in the lymphatic tissues, semen and CSF of CD8-depleted rhesus and cynomolgus macaques relative to nondepleted animals, implying the importance of CD8+ lymphocytes in limiting ZIKV dissemination and/or persistence in tissues. Lymphatic tissue viral loads were also consistently higher in cynomolgus compared to rhesus macaques, possibly reflecting the abbreviated time of infection before necropsy. Despite the detection of viral RNA in semen, ZIKV was scarce in reproductive tissues, in line with an absence of gross pathological lesions including atrophy of the testes. Although testicular atrophy is reported is murine models of ZIKV (Uraki et al., 2017), such manifestations have not been observed in macaques or in clinical cases (Matusali et al., 2018). In rhesus monkeys, it is possible that viral RNA in neural and reproductive tissues might have been only transiently present due to viral clearance given the 30-day infection period of these animals. Previous reports in rhesus (Aid et al., 2017) and pigtail (O’Connor et al., 2018) monkeys have also shown that ZIKV persists in lymphatic tissues well beyond the clearance of virus from the serum. It remains unclear whether the ZIKV present in lymph nodes is replication competent, but our data are nonetheless consistent with a model where the absence of CD8+ lymphocytes permits the dispersal of ZIKV.

CD8-depleted rhesus macaques also presented gross neural lesions at necropsy not seen in nondepleted animals. The most severe lesion occurred in the brainstem of a depleted animal that never recovered CD8+ lymphocytes, and similar manifestations of encephalomalacia and axon degeneration have been reported in ZIKV infection of human fetal brain tissue (Driggers et al., 2016; Petribu et al., 2018; Vesnaver et al., 2017). Perhaps complementarily, neural lesions in the CD8-recovering rhesus macaque were less severe. Although it is tempting to speculate that the absence of CD8+ lymphocytes in R25671 (CD8-depleted) and R64357 (CD8-depleted) allowed neural dissemination of the virus and thereby promoted neuropathy, our inability to detect ZIKV RNA in brain sections from these animals precludes this conclusion. Because ZIKV was cleared from the CSF of rhesus monkeys within 15 dpi, it is possible that virus could have also cleared from the CNS by necropsy and that these lesions were virus associated even if viral RNA was not detectable late in infection. Supporting this argument, CSF viral loads appear to be associated with ZIKV dissemination in neural tissue, given that the mock-depleted cynomolgus macaque C84545 showed the highest and most persistent CSF viremia and was also the only animal with ZIKV RNA identified in the brain. Despite the presence of CD8+ lymphocytes in C84545, this animal was the sole example of neural dissemination and occasionally produced responses more similar to CD8-depleted animals in key immune measures including NLR and classical monocyte frequency, underscoring the importance of innate immunity in limiting viral dissemination. Although it remains possible that off-target antibody effects produced immunological nuances in C84545, this animal ultimately aligned more closely with nondepleted animals in terms of serum viremia, antiviral gene induction, and adaptive immune activation, affirming the importance of CD8+ lymphocytes in maintaining immune regulation during acute ZIKV infection. The general absence of immune surveillance and IFN signaling in CD8-depleted animals might have permitted initial infection of neural tissues, which might have been transient due to the eventual priming of compensatory adaptive responses. Additionally, ZIKV localizes as discrete foci in macaque tissues (Hirsch et al., 2018), complicating the detection of sparse viral lesions within organs. It is worth noting that CNS localization of ZIKV has been observed as early as 5 dpi in acutely infected macaques (Osuna et al., 2016b), and a separate study in rhesus monkeys failed to identify ZIKV RNA in the CNS at 14 dpi, despite diffuse patterns of viral dissemination (Coffey et al., 2017). These findings, together with our own, establish precedence for early CNS dissemination of ZIKV in nonhuman primates, which may be cleared later in infection.

In summary, the present study illustrates a pliable dynamic between ZIKV and its hosts. CD8 depletion appears to alter innate immune activation and antiviral signaling and also modulate viral kinetics without overtly affecting serum viremia. Together with apparently compensatory adaptive responses and the presence of enhanced viral tissue distribution, these findings suggest that CD8 T cells provide default adaptive immune responses to ZIKV, a conclusion with important consequences for immune-based interventions such as vaccine development.

## Acknowledgements

This work was supported by a pilot award to N.J.M. from the TNPRC base grant (National Institutes of Health P51OD011104). The funders had no role in study design and interpretation.

## Author contributions

BS, AP, and NM planned the studies. BS, MF, ES, MW, KH, DS, RB, MG, LDM, and VD conducted the experiments. DW and MB provided reagents. BS, MF, MW, RB, AP, and NM interpreted the studies. BS and NM wrote the first draft. AP and NM obtained funding. All authors reviewed, edited, and approved the paper.

## Contribution to the Field

A number of reports have indicated an important role for CD8 T cells in the control of ZIKV replication in mice, but these models are limited in that the mice need to be immunocompromised for efficient viral infection. Using two established nonhuman primate models of ZIKV infection, we found that CD8+ lymphocytes are critical in orchestrating the earliest immune events during viral infection. The absence of CD8 cells enhanced viral dissemination into multiple tissues and prompted immediately observable host responses that diverged from previously consistent patterns of viremia and immunity. Importantly, the implications of these data may reach beyond ZIKV and are likely instructive to how CD8 cells interact with other immune cells to coordinate the control of viral infections generally.

## Materials & Correspondence

Material requests and correspondence should be addressed to N.J.M.

---

**Figure S1**

MT807R1 depletes CD8+ lymphocytes with variable recovery. **(A)** Absolute CD8 T cell counts in blood, as determined by complete blood count (CBC). **(B-C)** Absolute counts of CD4+/CD8+ double-positive T cells (B) and (C) CD4 T cells prior to infection, determined by CBC.

**Figure S2**

Comparison of virus and immune cell dynamics to a previous female cohort. Data from a previous cohort of ZIKV-infected non-pregnant female rhesus macaques is shown in gray, and data from rhesus macaques of the present study is overlaid. (**A**) Serum viral loads over the course of the studies (error bars, SD). (**B**) CSF viral loads over the course of the studies. (**C-D**) Frequencies of neutrophils (C) and total lymphocytes (D) in whole blood over time, determined by CBC. **(E)** NLR, derived using total neutrophil and lymphocyte CBC data.

**Figure S3**

Transcriptional responses correlate with serum viremia in nondepleted macaques. **(A-B)** Pathway analysis of gene expression in whole blood at 3 dpi relative to pre-infection reveals the induction of genes related to leukocyte homing (A) as well as differentially activated biological functions and disease-related pathways among CD8-depleted and non-depleted animals (B). **(C-D)** Patterns of antiviral gene induction at 1, 3, and 15 dpi in the whole blood of nondepleted (C) and CD8-depleted (D) rhesus macaques relative to pre-infection.

**Figure S4**

CD8 depletion modulates monocyte phenotype during ZIKV infection. (**A-C**) Flow cytometric analysis of monocyte activation, as measured by CD169 expression in classical (A), intermediate (B), and nonclassical (C) subsets in rhesus and cynomolgus macaques. (**D**) Overall monocyte activation, as measured by CD69 expression. (**E-F**) Expression of CD95 in classical (E) and nonclassical (F) subsets.

**Figure S5**

Reciprocal T cell responses are polyphenotypic. (**A-E**) Immunophenotyping of CD8 T cells in rhesus and cynomolgus macaques, including EM CD8 activation (A), CM CD8 activation (B) and proliferation (C), and naïve CD8 activation (D) and proliferation (E). **(F-J)** Immunophenotyping of CD4 T cells, including EM CD4 activation (F), CM CD4 activation (G) and proliferation (H), and naïve CD4 activation (I) and proliferation (J).

