## Supplemental Figures for "CD8+ lymphocytes modulate Zika virus dynamics and tissue dissemination and orchestrate antiviral immunity"

### Fig S1: MT807R1 depletes CD8+ lymphocytes with variable recovery

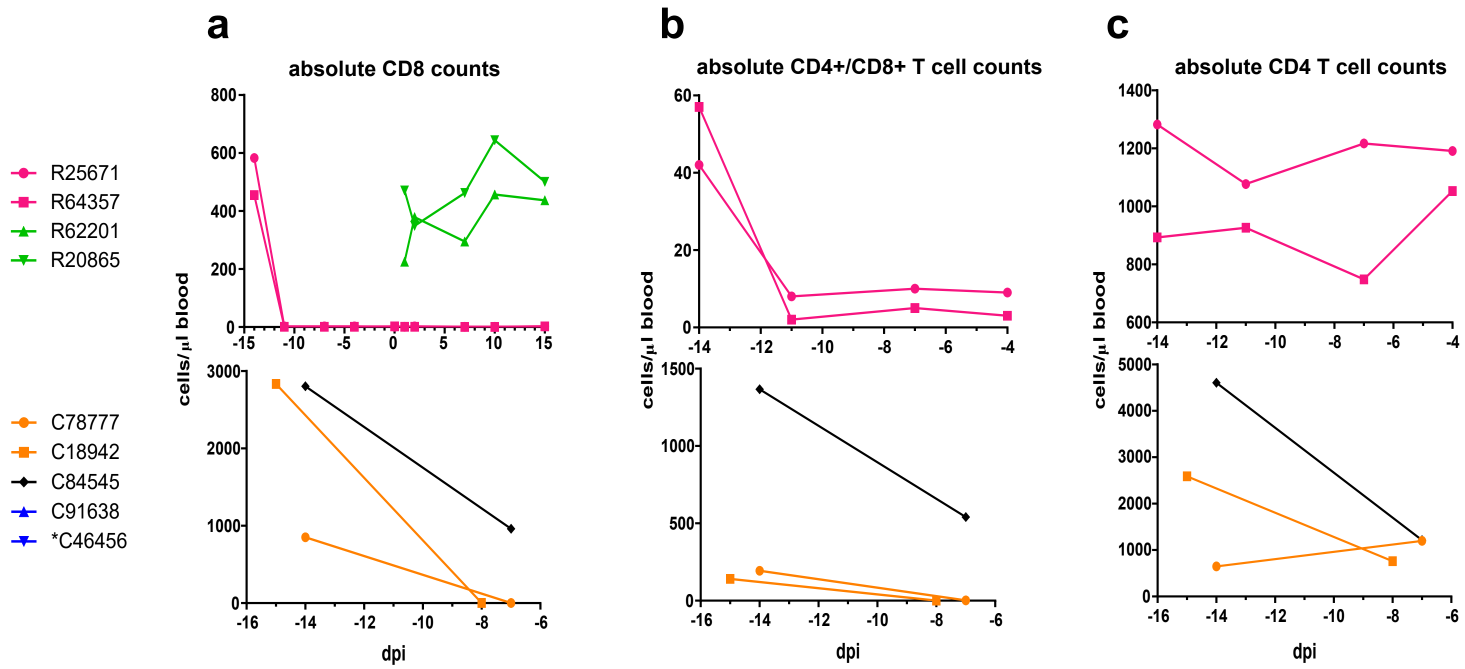

### Fig S2: Comparison of virus and immune cell dynamics to a previous female cohort

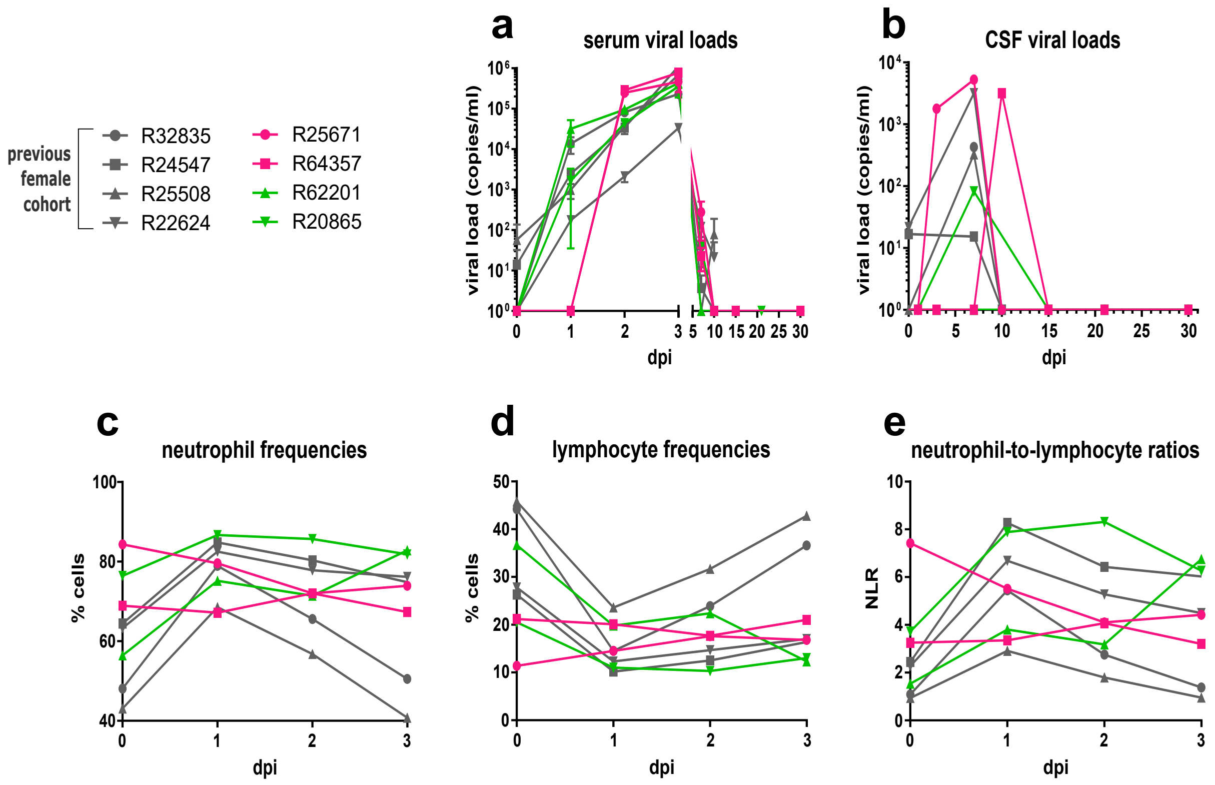

### Fig S3: Transcriptional responses correlate with serum serum viremia in nondepleted macaques

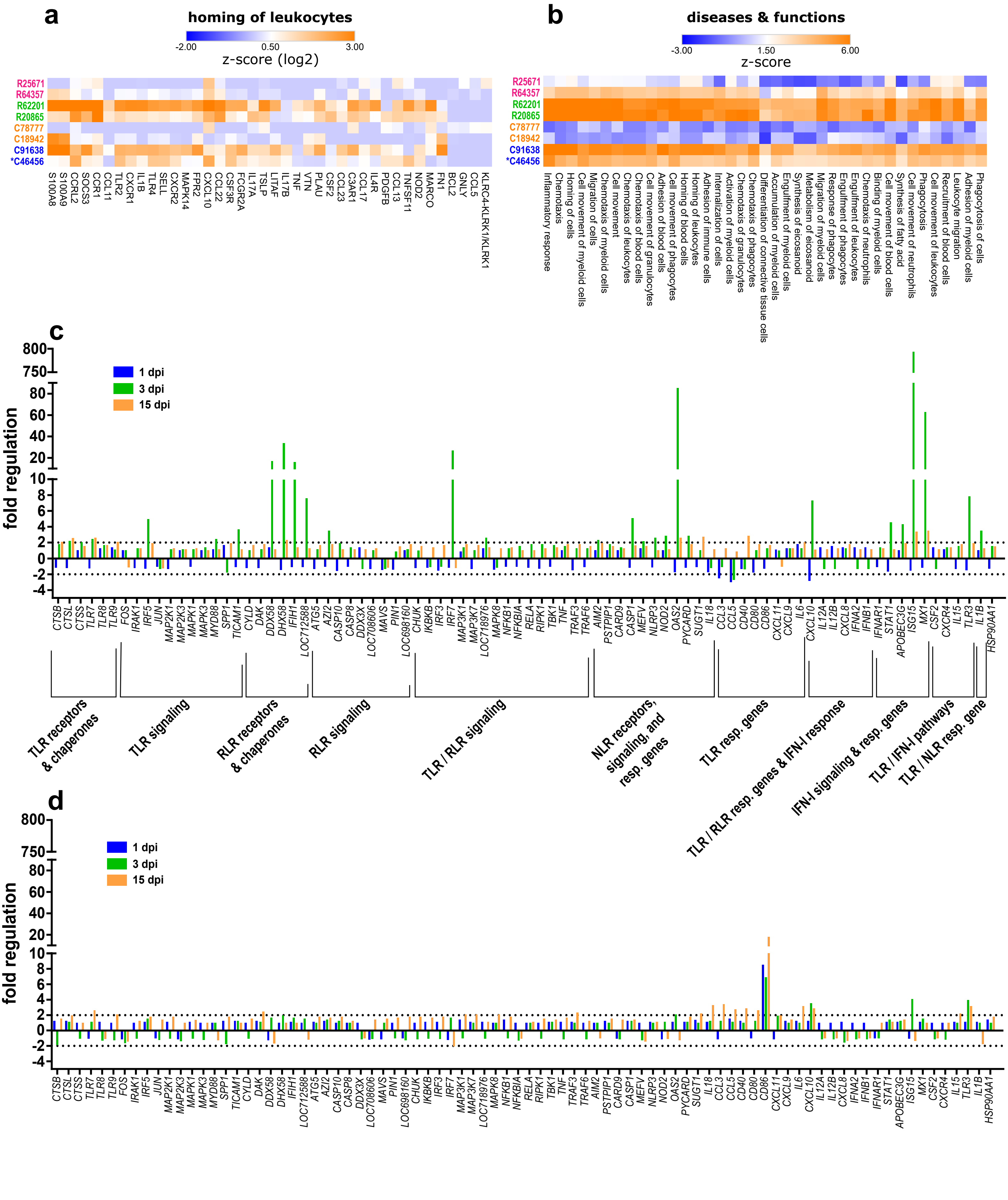

### Fig S4: CD8 depletion modulates monocyte phenotype during ZIKV infection

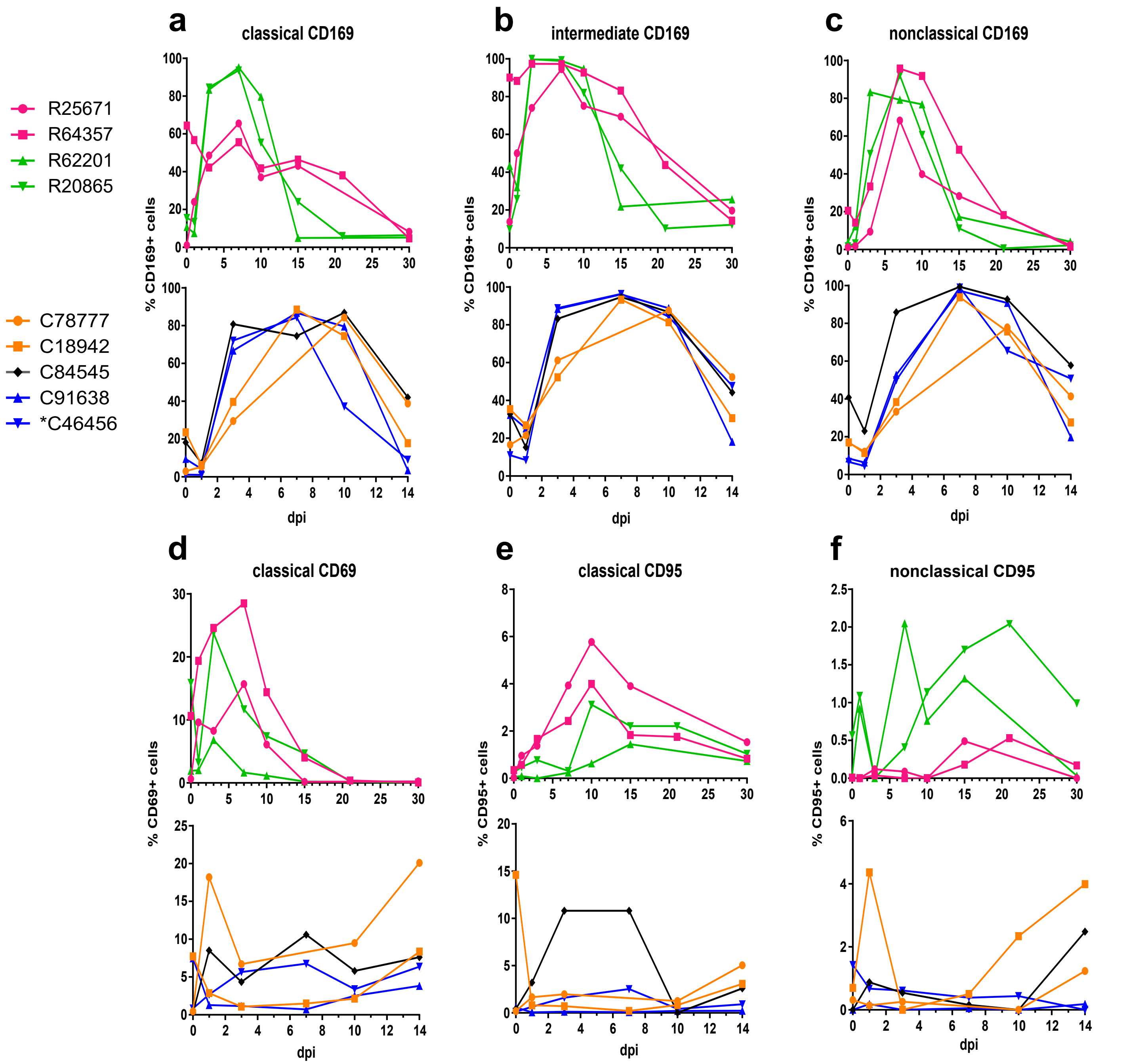

### Fig S5: Reciprocal T cell responses are polyphenotypic

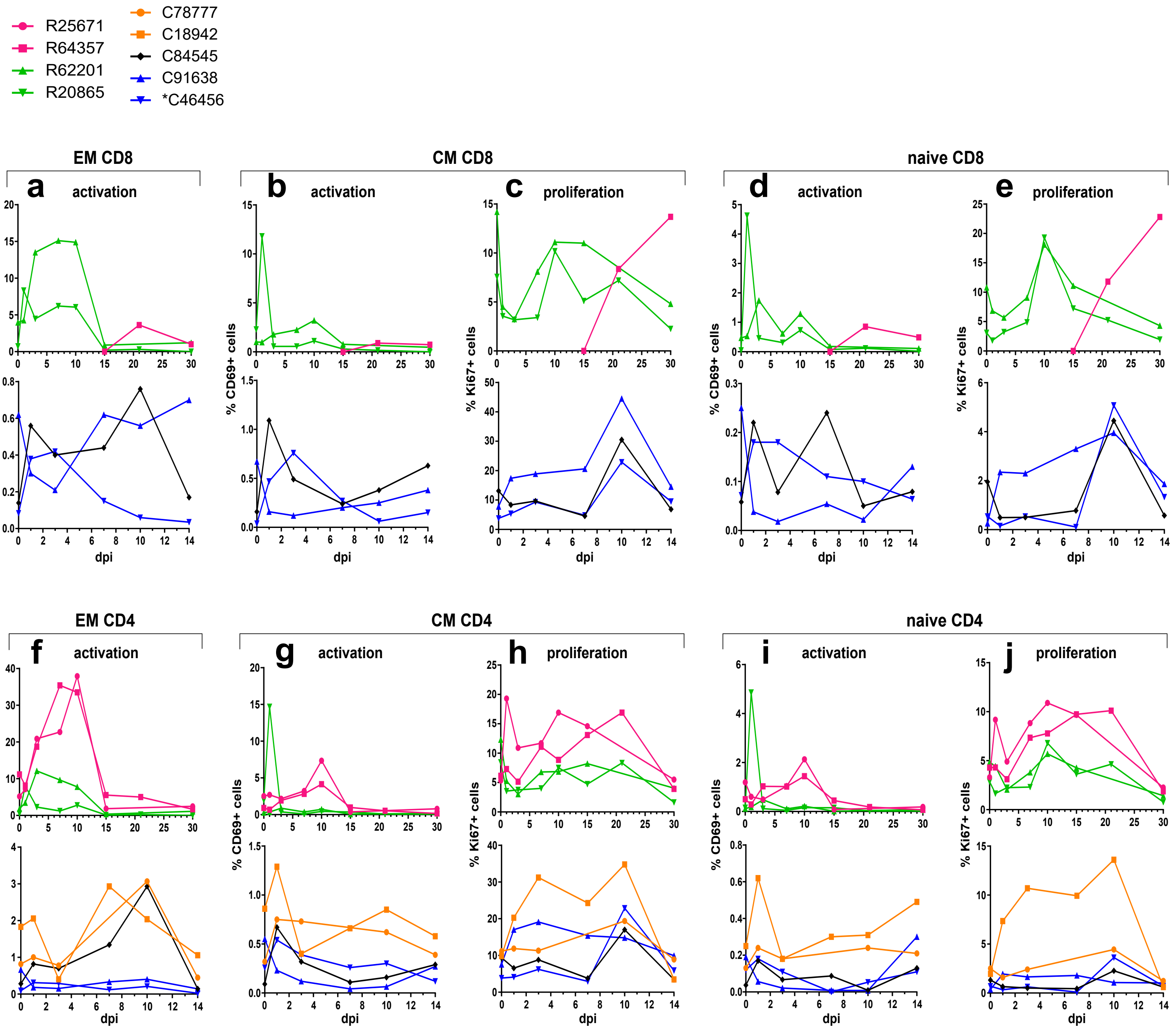
